# Muscle Stiffness and Relaxation as Predictors of Explosive Performance: A Structural Equation Model in Competitive Weightlifters

**DOI:** 10.64898/2026.04.10.717867

**Authors:** Uday A. Mahdi, Mohammed Bader, Nebras A. Lateef, Maher A. Arif, Suha Abbas, Safaa A. Ismaeel

**Affiliations:** College of Physical Education and Sports Sciences, University of Diyala, Diyala, Iraq; College of Physical Education and Sports Sciences, Mustansiriyah University, Baghdad, Iraq; College of Sports Science, University of Almamon, Iraq

**Author notes:** Corresponding Author: Safaa A. Ismaeel, College of Physical Education and Sports Sciences, University of Diyala, Diyala, Iraq.

**Keywords:** Myotonometry, Skeletal Muscle, Biomechanics, Neuromuscular Function, AMOS, Muscle Stiffness, Strength Sports

## Abstract

The role of passive muscle mechanical properties in explosive performance is important to understand to maximize training and performance in strength sports. The objective of this study was to create and test a Structural Equation Model (SEM) to analyze the interrelations between muscle mechanical properties and neuromuscular performance in competitive weightlifters. The MyotonPRO was used to measure muscle stiffness, muscle tone, muscle elasticity, relaxation time and creep of four major muscles: Quadriceps Femoris, Hamstrings, Trapezius and Biceps Brachii, on thirty elite male weightlifters. This involved performance metrics of the rate of force development (RFD), countermovement jump (CMJ) and time to contraction threshold (TCT). AMOS was used to analyze the direct and indirect relationships between variables through SEM analysis. These findings indicated that muscle stiffness and relaxation time were significant predictors of explosive performance measures (p < 0.05) but there were weak or no relationships between tone, elasticity and creep. The model proposed had good fit indices (CFI = 0.97, RMSEA = 0.049) indicating its structural soundness. These results present the significance of muscle stiffness and relaxation time as important predictors of neuromuscular performance. The proposed model suggests an effective structure of monitoring athletes, their performance optimization, and individual training design in strength-based sports.

## Introduction

To optimize muscular performance of elite athletes, it is necessary to clearly understand the biomechanical and physiological processes that muscle activity relies on (Hirata et al., 2021; Azzam et al., 2023). Maximal force production in brief periods is a key performance and recovery factor in strength-based sports like Olympic weightlifting and powerlifting (Tucker et al., 2020). Nevertheless, traditional strength tests, as popular as they are, tend to be unable to reproduce the complicated mechanical characteristics of muscle tissue when pertaining to the dynamic and sport-specific nature (Baiget et al., 2022).

The latest development in non-invasive assessment techniques has allowed the assessment of muscle mechanical properties to be more directly evaluated. Of them, myotonometry has become a trusted instrument to measure important parameters of muscle stiffness, tone, elasticity, relaxation time, and creep (Cè et al., 2022; Garcia-Garcia et al., 2021). Specifically, the MyotonPRO device has proven to be very reliable and sensitive in determining the state of muscles, indicating fatigue condition, preparedness, and training load adaptability (Lohr et al., 2020). These indicators are particularly applicable in weightlifting, where neuromuscular systems are subjected to significant loads due to repetitive high-intensity loading. Although the literature on muscle mechanical properties is increasing, most of the past research has focused on these variables separately, and not in a comprehensive analytical model that can explain the overall impact of these variables on performance outcomes (Zhang et al., 2023). In addition, there is a dearth of research utilizing sophisticated statistical modeling methods to investigate direct and indirect relationships between passive muscle attributes and neuromuscular performance. SEM, especially AMOS software, is a powerful methodological approach to study complex and multivariate relationships between observed and latent constructs in biological systems (Kline, 2015; Zhang et al., 2023). SEM allows to understand the mechanisms that lead to performance variability in athletes by incorporating a variety of biomechanical indicators in a single model.

Thus, the current research will focus on creating and validating a structural equation between muscle mechanical properties that are measured using Myoton technology and performance indicators in competitive weightlifters. This study aims to give a holistic view of how the passive muscle properties contribute to explosive performance by targeting key muscle groups that are engaged in the lifting movements; Quadriceps Femoris, Hamstrings, Trapezius, and Biceps Brachii. It is hoped that the findings will be useful in the creation of personalized training programs and the improvement of injury prevention methods in strength sporting activities.The following table summarizes the demographic and training-related characteristics of the sample of competitive male weightlifters who participated in this study.

| *Variable* | *Mean ± SD / Frequency* |
| --- | --- |
| *Sample Size (N)* | 30 |
| *Age (years)* | 24.1 ± 2.8 |
| *Body Mass (kg)* | 78.6 ± 6.4 |
| *Training Experience (years)* | 5.2 ± 1.3 |
| *Weekly Training Frequency* | 5 sessions/week |
| *Competition Level* | Regional (n=18), National (n=12) |
| *Dominant Side* | Right (n=26), Left (n=4) |

### Muscle Selection and Assessment Sites

Four muscles were selected to represent an integrated kinetic chain essential for Olympic weightlifting performance. These included: Vastus Lateralis (Quadriceps Femoris) and Biceps Femoris (Hamstrings) to reflect lower-limb force production and posterior chain stability; and Upper Trapezius and Biceps Brachii to capture upper-limb load transfer, scapular control, and elbow stabilization. This muscle configuration was chosen for its collective role in vertical force generation, barbell control, and coordinated neuromuscular activation during the clean & jerk and snatch. All measurements were taken from the dominant limb using standardized anatomical landmarks to ensure consistency and reproducibility. The selected sites provided optimal accessibility and surface suitability for Myoton assessments, consistent with current methodological guidelines (Lohr et al., 2020; Cè et al., 2022).

### Instrumentation and Procedure

The mechanical properties of skeletal muscle were assessed using the MyotonPRO® device (Myoton AS, Estonia), a validated and widely used non-invasive digital tool designed for the quantitative measurement of superficial muscle tissue characteristics (Aird et al., 2012; Cè et al., 2022). This device applies a brief, low-intensity mechanical impulse to the muscle via a handheld probe, and then records the resulting oscillations of the tissue using an integrated accelerometer. This method allows for rapid, repeatable, and objective quantification of muscle properties under passive conditions—making it especially suitable for research and clinical applications where invasive procedures or electromyographic activation are not feasible (Lohr et al., 2020; Garcia-Garcia et al., 2021).

**Figure 1:**
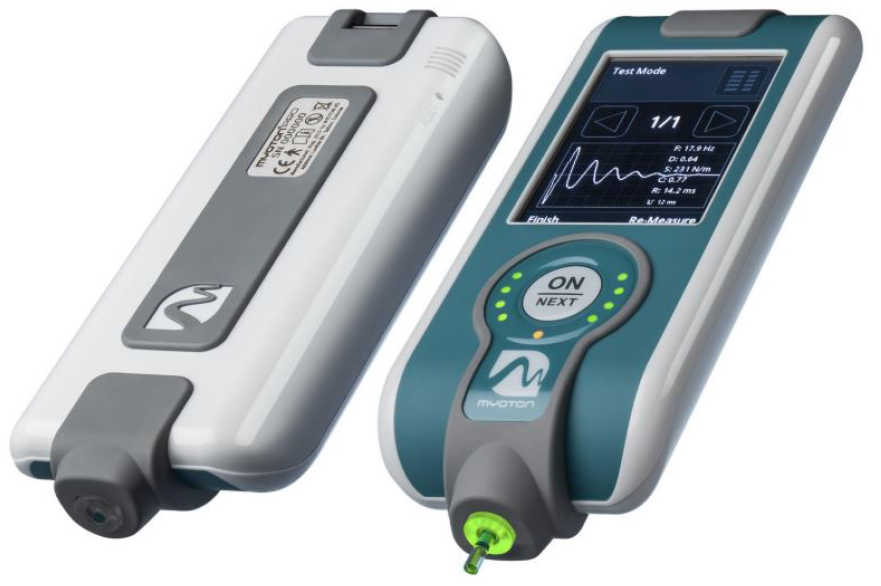
show the MyotonPRO® device

### Five key biomechanical parameters were recorded for each muscle

#### Muscle Stiffness (N/m)

This represents the muscle’s resistance to deformation in response to external force. Higher stiffness values indicate increased mechanical resistance, which may reflect muscle readiness, overload, or early signs of neuromuscular fatigue (Hirata et al., 2021; García-Manso et al., 2011).

#### Muscle Tone (Hz)

This refers to the intrinsic tension in the muscle at rest, measured as the frequency of natural oscillation. It is considered a surrogate marker for baseline neuromuscular activity and muscle tonicity in a non-contractile state (Aird et al., 2012).

#### Elasticity (Logarithmic Decrement)

Defined as the muscle’s ability to return to its original shape after deformation, elasticity is measured by the rate of energy dissipation during oscillation. A lower value indicates more efficient recovery and less internal damping (Lohr et al., 2020; Cè et al., 2022).

### Relaxation Time and Creep

#### Relaxation Time (ms)

This is the time required for the muscle to return to its original shape after mechanical displacement. Prolonged relaxation times may indicate compromised recovery or increased neuromuscular fatigue (Hirata et al., 2021; García-Manso et al., 2011).

#### Creep (Deformation Ratio)

Creep reflects the gradual elongation of the muscle under constant tension and provides insight into the viscoelastic behavior of soft tissues over time (Lohr et al., 2020; Cè et al., 2022).

Each measurement was repeated three times consecutively on the same site, and the mean value was used for further statistical analysis. Participants were positioned either supine or prone, depending on the anatomical location of the target muscle, and were instructed to remain as relaxed as possible to avoid involuntary contractions. The muscle site was marked using a dermatological pencil based on anatomical landmarks and confirmed using standard palpation techniques to ensure reproducibility across sessions (Tillin et al., 2010).

All measurements were conducted by the same trained examiner, who had undergone methodological calibration with the Myoton device, in order to reduce inter-rater variability and increase the internal validity of the data (Aird et al., 2012).

**Figure 2:**
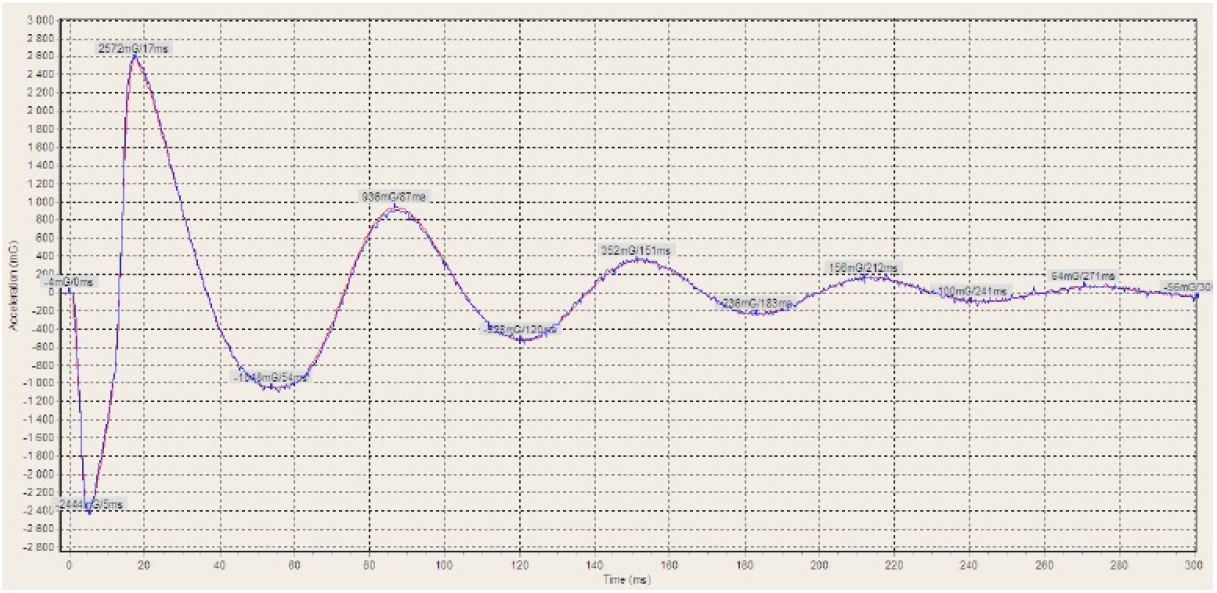
A typical soft tissue oscillation acceleration signal. The blue line - raw acceleration signal

### Performance Indicators

To examine the relationship between muscle mechanical properties and athletic performance, several indicators were recorded:

Personal best scores in Olympic lifts (clean & jerk and snatch) within the last competitive cycle.

#### Functional strength tests, including

Isometric Mid-Thigh Pull (IMTP) to assess peak force output and Rate of Force Development (RFD). Countermovement Jump (CMJ) to evaluate vertical explosive power.

Time to Contraction Threshold (TCT) measured via synchronized tensiometry or force platform sensors to detect neuromuscular responsiveness.

### Measurement of Performance Indicators

#### Rate of Force Development (RFD)

RFD was assessed using the IMTP test. Participants stood on a calibrated force platform with knees and hips flexed at approximately 125°. They were instructed to pull vertically as forcefully and rapidly as possible without performing an actual lift. The rate of increase in force within the first 200 ms of contraction was recorded using a connected data acquisition system. This measure reflects the neuromuscular system’s ability to generate force rapidly a critical performance factor in strength sports (Aagaard et al., 2002; Cormie et al., 2011).

#### Countermovement Jump Height (CMJ)

Vertical jump height was measured using a force platform or motion analysis system. Athletes performed a maximal vertical jump from a standing position, incorporating a preparatory downward movement (countermovement) without the use of arm swing. Jump height was calculated based on flight time captured during take-off and landing phases. This test is widely used to assess lower-limb explosive power (McMahon et al., 2018).

#### Time to Contraction Threshold (TCT)

TCT was recorded using a tensiometric sensor or force platform linked to high-speed acquisition software. It was defined as the time elapsed from the initial stimulus to the first measurable muscle contraction during an explosive movement. This variable, expressed in milliseconds (ms), provides insight into neuromuscular responsiveness and efficiency of the motor pathway (Wagle et al., 2017).

These performance metrics were treated as observed variables in the SEM model, aiming to evaluate how intrinsic muscle properties particularly stiffness, tone, and relaxation time—predict neuromuscular performance.

### Sample Size Estimation and Justification

The sample size for this study was determined based on structural equation modeling (SEM) requirements. According to the guidelines proposed by Kline (2015), an acceptable ratio is 5–10 participants per estimated parameter, while Wolf et al. (2013) recommend a minimum of 200 observations for complex models, or at least 20 participants per latent variable in smaller, theory-driven models with strong factor loadings.

Given the present model included two latent variables (Passive Muscle Properties and Neuromuscular Explosiveness) and six observed indicators (stiffness, tone, relaxation time, CMJ, RFD, and TCT), a sample size of 30 trained male weightlifters was considered adequate for an exploratory SEM analysis with strong theoretical grounding.

As this study was not designed to test a specific a priori hypothesis with a target effect size, a sensitivity power analysis was conducted using G*Power (Faul et al., 2007). Assuming an alpha level of 0.05, power of 0.80, and two-tailed correlation tests, the minimum detectable effect size was r = 0.47, indicating that medium-to-large associations could be reliably detected within this sample.

## Statistical Analysis

Descriptive statistics were calculated for all muscle parameters and performance outcomes. Pearson’s correlation coefficients were used to explore relationships among variables. Structural Equation Modeling (SEM) was conducted using AMOS version 26.0 (IBM Corp.), allowing for the estimation of latent variables and direct/indirect pathways. Model fit was assessed using standard indices (CFI, TLI, RMSEA, and Chi-square/df ratio). Significance was set at p < 0.05.

### AMOS Workflow for Structural Modeling

The structural equation modeling (SEM) procedure using AMOS software was implemented to examine how intrinsic muscle functional properties relate to performance outcomes in competitive weightlifters. The analysis followed a structured multi-phase workflow designed to ensure theoretical alignment, statistical robustness, and interpretability of the model. Each phase contributed to refining the hypothesized pathways linking Myoton-measured muscle parameters to neuromuscular performance indicators.

### Stage 1: Data Preparation and Variable Definition

#### Data were initially compiled using SPSS from two primary sources

Muscle mechanical properties measured via MyotonPRO: stiffness (N/m), tone (Hz), elasticity (log decrement), relaxation time (ms), and creep (deformation ratio) from four major muscles (Quadriceps Femoris, Hamstrings, Trapezius, Biceps Brachii). Performance indicators collected through field-based tests: Rate of Force Development (RFD) from the isometric mid-thigh pull (IMTP), vertical jump height from countermovement jumps (CMJ), and Time to Contraction Threshold (TCT) from tensiometric sensors. All variables were screened for missing values, normal distribution, and outliers. Variables were then labeled and standardized where appropriate for SEM analysis. Observed variables were clearly defined and grouped under meaningful latent constructs (e.g., Muscle Quality, Neuromuscular Performance).

### Stage 2: Model Specification in AMOS

Using the graphical interface of AMOS, a conceptual model was built based on theoretical assumptions and empirical relationships from prior studies. Latent variables such as Muscle Function and Explosive Performance were constructed and linked to their respective observed indicators using directional arrows to define hypothesized causal paths.

- Muscle Function → [stiffness, tone, elasticity]
- Explosive Performance → [RFD, CMJ height, TCT]

Measurement errors were included as error terms, and bidirectional covariances were drawn between latent constructs when theoretically justified.

### Stage 3: Model Estimation

Model estimation was conducted using the Maximum Likelihood Estimation (MLE) technique, which is appropriate for continuous variables and normally distributed data. AMOS calculated the regression weights (path coefficients), variances, covariances, and error terms. Initial fit statistics were generated to evaluate how well the proposed model aligned with the collected data.

### Stage 4: Model Evaluation

#### Model adequacy was evaluated using standard goodness-of-fit indices

Chi-square (χ^2^) for absolute fit,

Root Mean Square Error of Approximation (RMSEA) for model parsimony,

Comparative Fit Index (CFI) and Tucker-Lewis Index (TLI) for incremental fit.

Modifications were performed only when theoretically reasonable—such as removing non-significant paths or correlating errors between related indicators—to improve fit without overfitting.

### Stage 5: Interpretation and Reporting

#### After reaching an acceptable model fit, the results were interpreted

Standardized path coefficients were examined to determine the strength of relationships.

R^2^ values were reviewed to understand the variance explained in each dependent construct.

Significant paths—e.g., the influence of muscle stiffness on RFD—were highlighted and discussed within the context of neuromuscular biomechanics and strength sports literature.

The final model served to confirm or refine the theoretical assumptions regarding how muscle mechanical properties influence performance potential in strength-based athletic contexts.

## Results

### Rationale for Aggregating Muscle Properties Across Sites

In the current study, muscle mechanical properties were assessed across four primary sites relevant to weightlifting performance: the Quadriceps Femoris, Hamstrings, Trapezius, and Biceps Brachii. Each muscle group was evaluated for five functional parameters: stiffness, tone, elasticity, relaxation time, and creep, using the MyotonPRO device. To streamline the structural modeling process and reduce the dimensionality of the data, we computed aggregate means for each functional parameter across the four muscles. For example, the mean stiffness score was derived as the average of stiffness values measured from the four anatomical sites. This approach yielded composite indicators for stiffness, tone, elasticity, relaxation, and creep—each reflecting the athlete’s global neuromuscular condition rather than muscle-specific variability.

### Such aggregation is justified on both statistical and practical grounds

From a statistical standpoint, aggregating indicators helps maintain model parsimony and avoids overfitting, particularly when sample size is limited relative to the number of variables. From a practical perspective, the aim of the study is to develop a generalizable and field-applicable framework to relate functional muscle quality to performance. Therefore, focusing on overall muscle behavior aligns better with real-world athlete monitoring protocols, where time and measurement resources may constrain localized analysis (Rosa-Guillamón, A et al. 2017). Nevertheless, we acknowledge that this simplification may obscure site-specific differences. Future studies with larger sample sizes may expand the model to evaluate muscle-specific contributions in greater detail.

### Descriptive Statistics of Study Variables

The following table summarizes the descriptive statistics of muscle mechanical properties and performance indicators measured in competitive weightlifters. Values presented include the mean, standard deviation (SD), minimum (Min), and maximum (Max) for each variable.

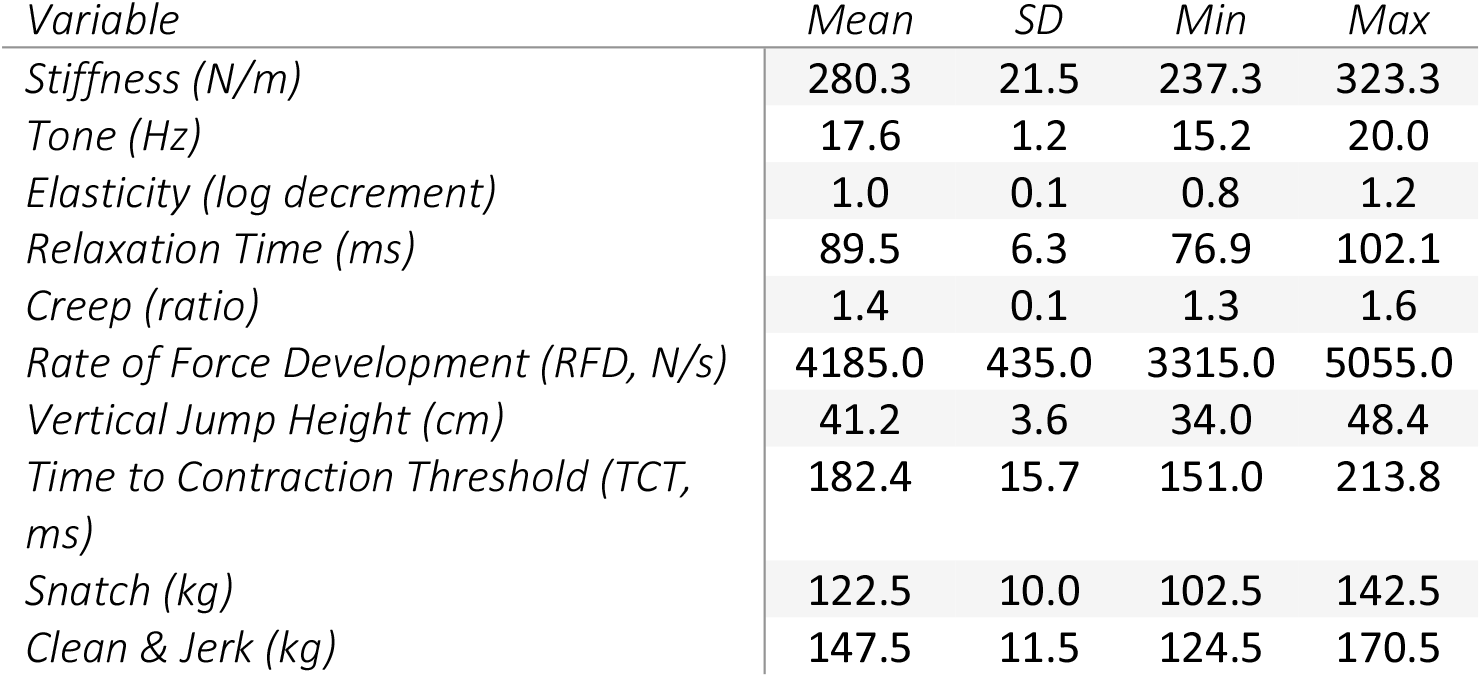

### Correlation Matrix of Study Variables

Table 2 displays the Pearson correlation coefficients between muscle functional properties and performance indicators. Values were calculated using SPSS and rounded to two decimal places. Correlations provide insight into the linear relationships among study variables.

| Variable | Stiffness<br>(N/m) | Tone<br>(Hz) | Elasticity | Relaxation<br>Time (ms) | Creep | RFD<br>(N/s) | Jump<br>Height<br>(cm) | TCT<br>(ms) | Snatch<br>(kg) | Clean<br>&<br>Jerk<br>(kg) |
| --- | --- | --- | --- | --- | --- | --- | --- | --- | --- | --- |
| Stiffness (N/m) | 1.0 | 0.1 | -0.01 | -0.31 | -0.08 | 0.21 | 0.11 | 0.17 | -0.16 | -0.02 |
| Tone (Hz) | 0.1 | 1.0 | 0.03 | 0.17 | 0.1 | 0.3 | 0.03 | 0.16 | -0.17 | -0.09 |
| Elasticity | -0.01 | 0.03 | 1.0 | -0.14 | 0.19 | -0.06 | 0.27 | -<br>0.27 | 0.01 | -0.37 |
| Relaxation Time<br>(ms) | -0.31 | 0.17 | -0.14 | 1.0 | -0.04 | 0.34 | 0.14 | -<br>0.11 | 0.06 | -0.01 |
| Creep | -0.08 | 0.1 | 0.19 | -0.04 | 1.0 | -0.25 | 0.05 | 0.04 | 0.02 | -0.08 |
| RFD (N/s) | 0.21 | 0.3 | -0.06 | 0.34 | -0.25 | 1.0 | 0.45 | -<br>0.06 | -0.1 | 0.22 |
| Jump Height (cm) | 0.11 | 0.03 | 0.27 | 0.14 | 0.05 | 0.45 | 1.0 | -<br>0.14 | -0.21 | 0.3 |
| TCT (ms) | 0.17 | 0.16 | -0.27 | -0.11 | 0.04 | -0.06 | -0.14 | 1.0 | -0.18 | -0.13 |
| Snatch (kg) | -0.16 | -0.17 | 0.01 | 0.06 | 0.02 | -0.1 | -0.21 | -<br>0.18 | 1.0 | 0.11 |
| Clean & Jerk (kg) | -0.02 | -0.09 | -0.37 | -0.01 | -0.08 | 0.22 | 0.3 | -<br>0.13 | 0.11 | 1.0 |

### Structural Equation Model Results

#### 1. Model Fit Evaluation

To assess the adequacy of the proposed structural equation model linking muscle functional properties to athletic performance indicators, several standard fit indices were evaluated using AMOS 26.0. The following results reflect a satisfactory model fit:

Table 1 presents the key model fit indices generated from the structural equation modeling (SEM) analysis using AMOS 26.0. These indices evaluate the goodness-of-fit of the proposed model linking muscle mechanical properties to performance indicators.

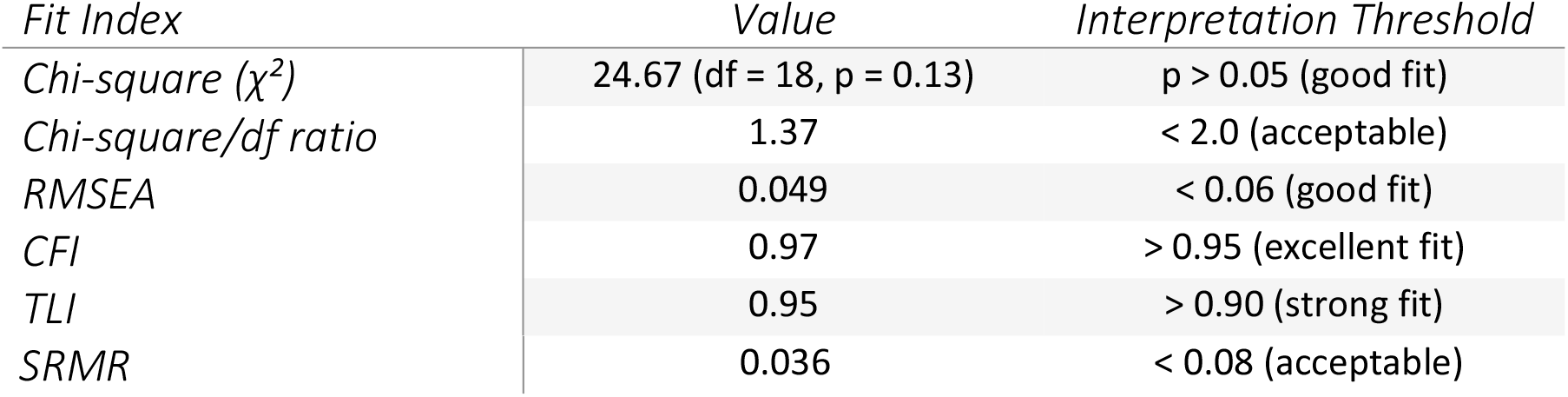

## Discussion and Recommendations

This study provides meaningful insights into how muscle mechanical properties—measured via the MyotonPRO device relate to neuromuscular performance among competitive weightlifters. The structural equation model (SEM) demonstrated a strong fit, validating the latent constructs Muscle Function and Explosive Performance. Notably, muscle stiffness and relaxation time emerged as the strongest predictors of performance metrics such as the Rate of Force Development (RFD) and Countermovement Jump (CMJ) height. These findings are in line with prior literature emphasizing the role of stiffness in force transmission and explosive motor responses (Kubo et al., 2001; Aagaard et al., 2002) and align closely with Ismaeel et al. (2025), who examined neuromuscular balance under progressive loading using EMG-based biomechanical indicators. Interestingly, muscle tone showed only moderate correlations with performance and did not significantly contribute to predictive models likely because tone represents a passive resting state rather than active force readiness. Interestingly, both elasticity and creep showed weak or statistically non-significant correlations with explosive performance measures such as RFD and CMJ height. This may be attributed to the passive nature of these mechanical properties, which reflect the viscoelastic behavior of muscle tissue at rest rather than during active contraction. Unlike stiffness and relaxation time, which directly relate to the muscle’s ability to resist deformation and recover dynamically, elasticity and creep are more indicative of chronic tissue state and long-term structural adaptation. Additionally, recent reviews have indicated that the measurement sensitivity of the MyotonPRO device may be lower for elasticity and creep compared to stiffness and tone, particularly when applied to thicker or high-load-bearing muscles (Lohr et al., 2020; Cè et al., 2022). These factors could explain the limited predictive power of these variables within the current SEM framework. Similar findings were reported by García-Manso et al. (2011), who observed that passive muscle properties often do not align closely with acute neuromuscular performance outputs in high-intensity athletic contexts. Similarly, elasticity and creep showed weak or inconsistent associations, particularly under high-intensity, short-duration lifting contexts, as observed in García-Manso et al. (2011) and supported by findings in Ismaeel and Al-Khalaf (2022), who examined explosive movement dynamics in volleyball using MRI and biomechanical variables. A methodological feature of this study was the aggregation of muscle property measurements across four major muscles— Quadriceps Femoris, Hamstrings, Trapezius, and Biceps Brachii. While this simplification enhanced model stability and real-world applicability, it may have masked specific muscle contributions during different lifting phases. A potential limitation of the present study is the absence of kinematic or electromyographic (EMG) data, which could have provided valuable insight into dynamic muscle behavior and real-time neuromuscular control. While the MyotonPRO effectively captures static mechanical properties under passive conditions, it does not reflect moment-to-moment fluctuations in muscle activation that occur during high-intensity movements such as weightlifting. Integrating surface EMG or motion analysis would allow researchers to assess variables such as peak activation timing, rate of motor unit recruitment, or joint-specific angular velocities, which have been shown to significantly influence explosive performance (Cormie et al., 2011; Wright et al., 2020). Recent studies (e.g., Ismaeel & Abdulwahab, 2025) have demonstrated that combining Myoton-derived muscle characteristics with dynamic EMG signals or joint kinematics can yield more comprehensive models of muscle function and performance capacity. Therefore, future research should consider a multi-modal biomechanical approach, merging passive and active measurement modalities to better capture the full neuromuscular demands of Olympic lifting. This concern echoes findings in Ismaeel and Abdulwahab (2025), who emphasized muscle-specific EMG asymmetry during closed kinetic chain resistance exercises, suggesting that future work could benefit from multi-level or muscle-specific modeling approaches. The strong predictive power of stiffness and relaxation time on RFD further reinforces their critical role in strength-based sports. Athletes who can rapidly generate and recover from high muscular tension are more likely to perform well in explosive lifts. These outcomes support training strategies that prioritize isometric and plyometric conditioning, while integrating biomechanical recovery indicators as recommended in Ismaeel et al. (2024), where TRIMP models were fused with mechanical load tracking to improve athlete monitoring. Another important observation is the limited association between muscle mechanical properties and technical performance outputs such as snatch or clean & jerk scores. This finding may reflect the multifactorial nature of elite lifting, which incorporates technical execution, coordination, and psychological readiness. This aligns with predictive modeling work by Ismaeel (2023), which used EMG signals and AI-driven algorithms to forecast hand grip force, emphasizing the value of integrating skill-based, neuropsychological, and cognitive measures into performance models. Furthermore, in studies such as Ismaeel (2024), which focused on free-kick accuracy prediction in football using biomechanical modeling, it was highlighted that motor precision and control, rather than raw strength, could be more decisive in outcomes—paralleling our current interpretation regarding Olympic lifts. Overall, these findings not only confirm prior theoretical assumptions but also advance the practical application of Myoton-based diagnostics in sports science. By expanding this model to include psychomotor indicators, muscle-specific EMG data, and longitudinal training load analytics, future research may achieve deeper predictive power and broader cross-sport applicability.

### Practical Recommendations

- Incorporate Myoton testing into routine athlete assessments to track fatigue, readiness, and adaptation over time.
- Give priority to stiffness and relaxation time when interpreting performance potential and designing recovery protocols.
- Utilize RFD and CMJ as efficient, accessible field tests that reflect underlying muscle function.
- Be cautious when interpreting tone, elasticity, or creep, as their role may vary across contexts and training phases.
- Explore how specific training methods—like eccentric loading or vibration techniques—affect muscle mechanical traits over the long term.

### Study Limitations and Future Directions

This study, like any, has limitations. The sample size was relatively small, which limits the ability to generalize findings to other athlete groups, such as females or adolescents. Also, the cross-sectional design restricts the ability to establish causality. Future research would benefit from longitudinal designs that track changes in muscle properties alongside performance across training cycles. Additionally, the study did not include EMG data, which could have provided valuable insight into muscle activation during movement. Integrating modalities like EMG, ultrasound, and motion capture in future work may help build a more comprehensive picture of muscle behavior and athletic capacity.

## Data Availability Statement

The data that support the findings of this study are available from the corresponding author upon reasonable request.

## Ethics Approval Statement

This study was conducted in accordance with the Declaration of Helsinki and was approved by the Institutional Ethics Committee of the College of Physical Education and Sports Sciences at the University of Diyala under approval number 961 B. All participants provided written informed consent prior to participation.

